# Nanotube-mediated cross-feeding couples the metabolism of interacting bacterial cells

**DOI:** 10.1101/114462

**Authors:** Shraddha Shitut, Tobias Ahsendorf, Samay Pande, Matthew Egbert, Christian Kost

## Abstract

Bacteria frequently engage in cross-feeding interactions that involve an exchange of metabolites with other micro- or macroorganisms. The often obligate nature of these associations, however, hampers manipulative experiments, thus limiting our mechanistic understanding of the ecophysiological consequences that result for the organisms involved. Here we address this issue by taking advantage of a well-characterised experimental model system, in which auxotrophic genotypes of *E. coli* derive essential amino acid from prototrophic donor cells using intercellular nanotubes. Surprisingly, donor-recipient cocultures revealed that the mere presence of auxotrophic genotypes in coculture was sufficient to increase amino acid production levels in donor cells. Subsequent experiments unravelled that this effect was due to the depletion of amino acid concentrations in the cytoplasm of donor cells, which delayed feedback inhibition of the corresponding amino acid biosynthetic pathway. This finding indicates that in newly established mutualistic associations, an intercellular regulation of exchanged metabolites can simply emerge from the architecture of the underlying biosynthetic pathways, rather than through the evolution of new regulatory mechanisms. Taken together, our results show that a single loss-of-function mutation can physiologically couple the metabolism of two cross-feeding cells in a source-sink-like relationship.

---

**Originality-Significance Statement**: The results of our study indicate that the distribution of metabolites within networks of interacting bacterial cells may be self-organized by local interactions among neighbouring cells rather than requiring a super-ordinated regulatory system.

## INTRODUCTION

Bacteria often engage in metabolic cross-feeding interactions with other bacteria and eukaryotic organisms (Kiers, Rousseau et al. 2003, Belenguer, Duncan et al. 2006, Vogel and Moran 2011, Johnson, Goldschmidt et al. 2012, McFall-Ngai 2014, Seth and Taga 2014, Ponomarova and Patil 2015, Zelezniak, Andrejev et al. 2015, Estrela, Kerr et al. 2016). In many of these cases, two or more interacting partners reciprocally exchange primary building block metabolites such as amino acids (Payne, Rouatt et al. 1957, Junglas, Briegel et al. 2008, Sieuwerts, Molenaar et al. 2010, Vogel and Moran 2011, Garcia, Buck et al. 2015), vitamins (Croft, Lawrence et al. 2005, Rodionova, Li et al. 2015), or even nucleotides (Sieuwerts, Molenaar et al. 2010, Dean, Hirt et al. 2016, Loera-Muro, Jacques et al. 2016). However, why do organisms produce these compounds to benefit others, rather than using these metabolites for themselves? Recent empirical and theoretical evidence suggests that adaptive benefits resulting from the loss of biosynthetic functions may drive the establishment of such metabolic interactions: Organisms may release metabolites into the extracellular environment, for example as a consequence of a leaky membrane (Shiio, Ocirc et al. 1962) or overflow metabolism (Paczia, Nilgen et al. 2012). Any mutant that has lost the ability to produce the corresponding compounds will start to use these environmentally available metabolite pools (D’Souza, Waschina et al. 2014, D’Souza and Kost 2016). Since auxotrophic mutants save the costs of producing the focal metabolites by themselves, they gain a selective advantage over other, prototrophic cells that still produce them (Morris, Lenski et al. 2012, D’Souza, Waschina et al. 2014). Moreover, in some species, auxotrophy-causing mutations even trigger the formation of intercellular nanotubes (Pande, Shitut et al. 2015). These are membrane-based structures that help auxotrophic bacteria to derive amino acids from the cytoplasm of other bacteria, thus enhancing metabolite transfer between cells.

The idea that adaptive gene loss causes the establishment of obligate cross-feeding interactions has been termed the *black queen hypothesis* (Morris, Lenski et al. 2012, Morris 2015). Indeed, finding that metabolic auxotrophies are crucially involved in many naturally-occurring cross-feeding interactions corroborates this interpretation (Croft, Lawrence et al. 2005, Giovannoni, Tripp et al. 2005, Sahu and Ray 2008, Garcia, Buck et al. 2015, Hubalek, Buck et al. 2017). Since gene loss seems to be a key step in the establishment of metabolic cross-feeding interactions, most naturally evolved systems are characterized by obligate dependencies among the interacting parties. This means that the participating genotypes can usually not be cultivated in isolation, thus impeding experimental manipulation (Pande and Kost 2017). As a consequence, mechanistic details on how the transition into a metabolic mutualism affects the physiology of the strains involved remain poorly understood. For example, it is not clear whether or not a unidirectional exchange of amino acids via nanotubes incurs fitness costs to donor cells? Moreover, it is unknown how the consumption of amino acids by a recipient affects amino acid production of a donor cell?

Here we address these questions taking advantage of a previously established model system, in which two genotypes of *E. coli* unidirectionally exchange essential amino acids. These one-way cross-feeding interactions were established by matching amino acid donors with auxotrophic recipients that obligately required the corresponding amino acid for growth. Utilizing genetically engineered single gene deletion mutants for this purpose ruled out pre-existing traits that arose as a consequence of a coevolutionary history among interaction partners. Moreover, a focus on unidirectional cross-feeding excluded confounding effects that may occur in reciprocal interactions such as for example self-enhancing feedback loops (Kun, Papp et al. 2008). Finally, taking advantage of intracellular reporter constructs allowed analysing both internal amino acid pools as well as their production levels in real-time under *in vivo* conditions.

Our experimental results revealed not only that auxotrophic recipients used nanotubes to derive amino acids from prototrophic cells, but also that this process increased the production of the focal amino acid in donor cells. This was due to a drop in amino acid-concentrations in the cytoplasm of donor cells, which delayed the feed-back inhibition of the corresponding biosynthetic pathway and in this way increased production levels of the focal amino acid. In other words, a nanotube-mediated exchange of cytoplasmic amino acids coupled the metabolism of two interacting partners in a source-sink-like relationship. These results show for the first time that regulatory mechanisms that control the production of amino acids in one cell, can easily be extended to include other cells as well, provided the interaction is based on an intercellular exchange of metabolites via nanotubes.

## RESULTS

### Construction and characterisation of unidirectional cross-feeding interactions

To establish unidirectional cross-feeding interactions within *Escherichia coli*, five different genotypes served as amino acid donors: Two single gene deletion mutants (Δ*mdh* and Δ*nuoN*) that produce increased amounts of several different amino acids (Pande, Merker et al. 2014), two deletion mutants that produce increased amounts of either histidine or tryptophan (Δ*hisL* and Δ*trpR*) (Pande, Kaftan et al. 2015), as well as unmanipulated *E. coli* WT cells (Figure 1, Supplementary Table 1). Three genotypes served as recipients, which were auxotrophic for the amino acids histidine (Δ*hisD*), lysine (Δ*lysR*), and tryptophan (Δ*trpB*) (Figure 1, Supplementary Table 1) and thus essentially required an external source of these metabolites to grow (Bertels, Merker et al. 2012).

**Figure 1.**
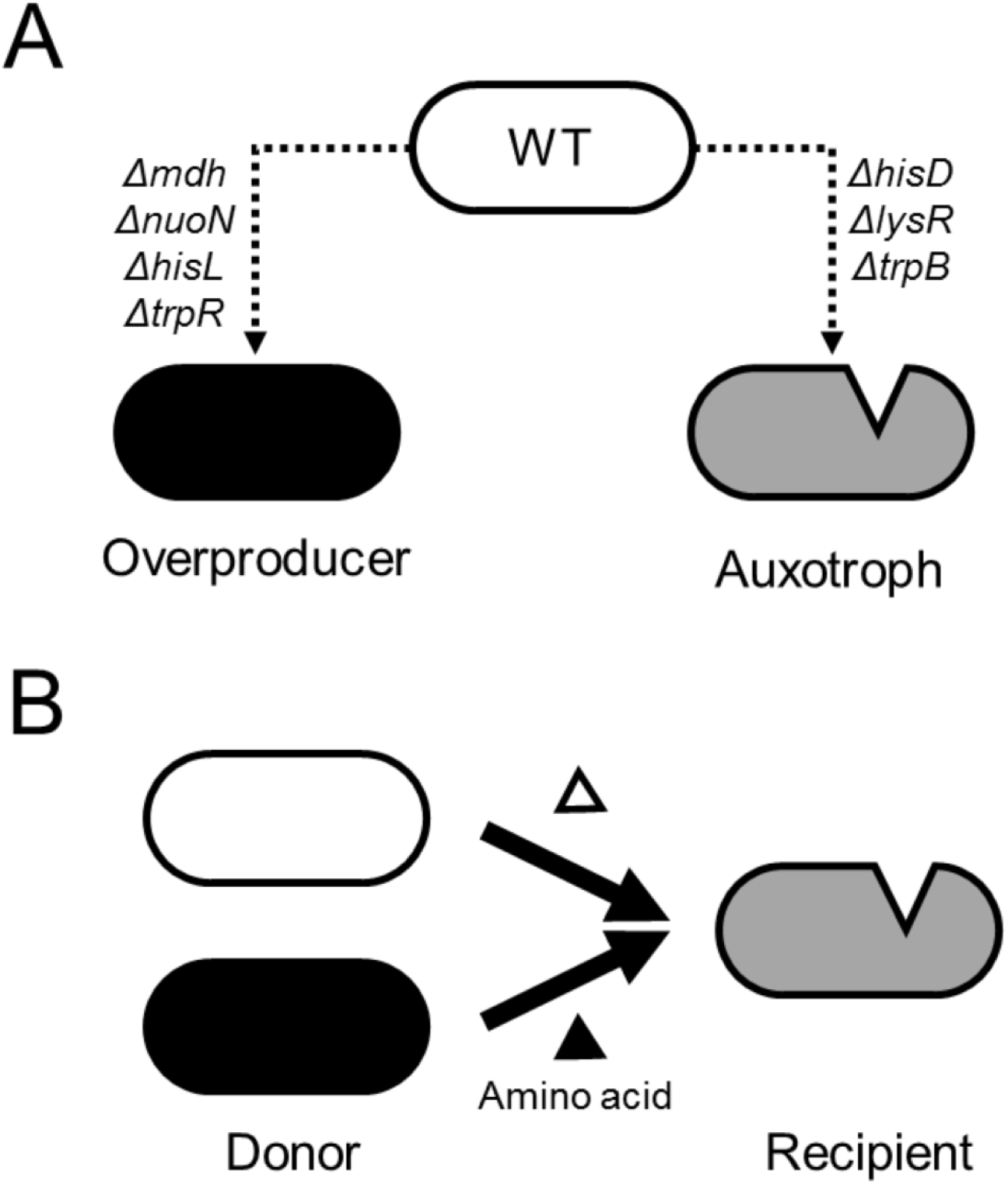
Experimental system used. (**A**) Design of genotypes. Single genes were deleted from *E. coli* BW25113 wild type (WT) to generate mutants that produce increased amounts of amino acids (overproducer) as well as mutants that essentially require a certain amino acid to grow (auxotroph). (**B**) Coculturing an amino acid donor (i.e. WT or overproducer) together with an auxotrophic recipient results in a one-way cross-feeding interaction that is obligate for the recipient, but not the donor.

As a first step, we quantified the amounts of amino acids the five donor strains produced in monoculture during 24 hours of growth. Analysing culture supernatant and cytoplasm of the focal donor populations using LC/MS/MS revealed that Δ*nuoN* produced significantly increased amounts of histidine, lysine, and tryptophan in both fractions relative to the WT (Mann Whitney U-test: P<0.05, n=4, Supplementary Figure 1), while production levels of the Δ*mdh* mutant did not differ significantly from WT-levels (Mann Whitney U-test: P>0.05, n=4, Supplementary Figure 1). Similarly, both the intra- and extracellular concentrations of tryptophan in the Δ*trpR* mutant were significantly elevated over WT-levels (Mann Whitney U-test: P<0.05, n=4, Supplementary Figure 1). In contrast, Δ*hisL* released twice as much of histidine into the growth medium as was released by the WT (two sample Mann Whitney test: P<0.05, n=4, Supplementary Figure 1), while it contained much lower levels of histidine in its cytoplasm than the WT.

### Intercellular transfer of amino acids is contact-dependent

Taking advantage of a set of well-characterised genotypes, we addressed the question whether donor and recipient cells exchange amino acids in coculture, and if so, whether this interaction is contact-dependent. To this end, populations of donor and recipient cells were cocultured in a device (i.e. *Nurmikko cell*), in which both partners can either be grown together in the same compartment or be separated by a filter membrane that allows passage of small molecules, yet prevents direct physical interactions between bacterial cells (Pande, Shitut et al. 2015). Inoculating donor and recipient strains in different combinations revealed, in all tested cases, growth of auxotrophic recipients when they were not physically separated from donors (Figure 2A-C). Auxotrophic recipients grew significantly better when cocultured with amino acid overproducers (Δ*mdh*, Δ*nuoN*, Δ*hisL*, and Δ*trpR*) than with the WT (Dunnett’s T3 post hoc test: P<0.05, n=4). However, introducing a filter membrane to physically separate donor and recipient cells effectively eliminated growth of recipients in all cases. Surprisingly, the process of coculturing in this way did not affect the growth of donor populations (Figure 2A-C). This observation suggested that even though donor cells had to produce all the amino acids required by auxotrophs for growth (i.e. His, Lys, and Trp), these increased production level did not significantly affect the final growth of donor genotypes. To further evaluate whether the observed increased amino acid production levels did not incur a fitness cost to donor cells, the growth rates of donor populations in mono- and coculture during the exponential phase were compared. This experiment revealed a cost of amino acid production that only affected overproducing genotypes (i.e. Δ*mdh*, Δ*nuoN*, and Δ*trpR*) when paired up with the tryptophan-auxotrophic recipient Δ*trpB* (Figure 2D). In these cases, the growth rate of donor genotypes in coculture was significantly reduced relative to their growth in monocultures (FDR-corrected paired sample t-tests: P<0.05, n=6). In all other cases, with WT as donor and lysine- or histidine auxotrophic recipients, no fitness cost was detected (FDR-corrected paired sample t-tests: P>0.05, n=6). Three main insights result from this experiment: First, even though producing the amino acids auxotrophs required for growth significantly reduced the growth rate of donors in some cases, this fitness cost was not detectable on the level of the population size after 24 h of growth in coculture. Second, the total productivity of donors and recipients was significantly increased when cells were cocultured as compared to the situation when they were physically separated by a filter membrane (Mann Whitney U-test: P<0.05, n=4, Figure 2A-C). Third, physical contact between donor and recipient cells was required for a transfer of amino acids between cells.

**Figure 2.**
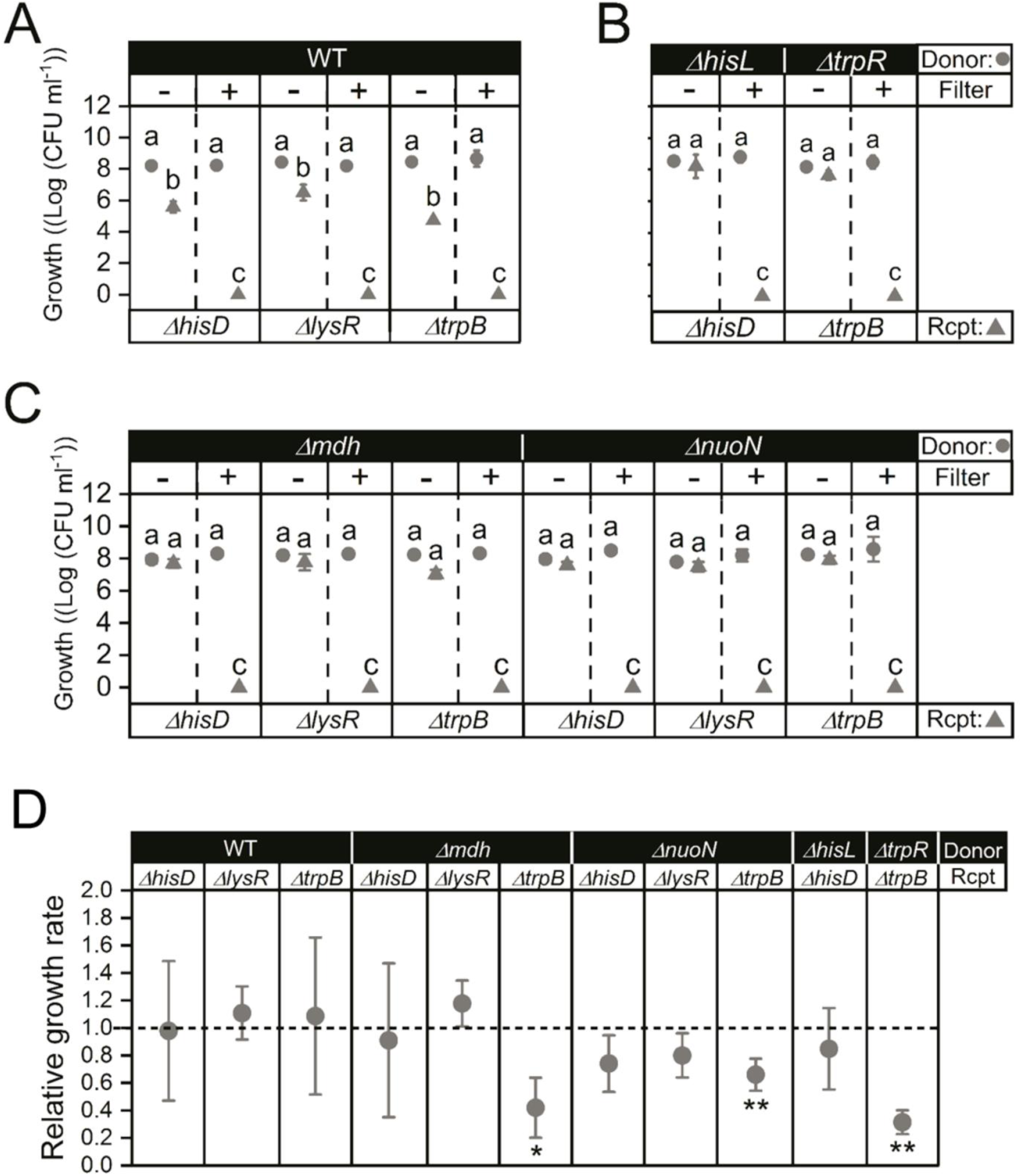
Contact-dependent exchange of cytoplasmic amino acids. (**A-C**) Amino acid exchange is contact-dependent. Amino acid donors (circles) were cocultured with auxotrophic recipients (Rcpt, triangles); Δ*hisD*, Δ*lysR*, Δ*trpB*. Strains were cultured either together in the same compartment (− Filter) or separated by a filter membrane (+ Filter) that allows passage of free amino acids, but prevents direct physical contact between cells. Growth over 24 h was determined as number of colony-forming units (CFU) per ml by subtracting the value at 0 h from that reached at 24 h. Different letters indicate significant differences (Dunnett’s T3 post hoc test: P<0.05, n=4). (**D**) Cost of amino acid overproduction. The growth rate of the donors (WT and overproducers) when grown in coculture with auxotrophic recipients is plotted relative to that in monocultures (dashed line). Growth rate was calculated as slope of the growth curve over the exponential phase (14 h to 20 h after inoculation). Asterisks indicate significant differences of growth rates between mono- and cocultures (paired t-test: ** P< 0.01, * P<0.05, n=4). In all cases, mean (±95% confidence interval) are shown.

### Cytoplasmic constituents are transferred from donor to recipient cells via nanotubes

The observation that metabolite cross-feeding among cells was contact-dependent suggested that separating cells with a physical barrier prevented the establishment of structures required for amino acid exchange. A possible explanation for this could be intercellular nanotubes, which would allow direct transfer of cytoplasmic amino acids from donor to recipient cells (Pande, Shitut et al. 2015). Indeed, microscopic analysis revealed the presence of nanotubes connecting cells in a coculture of donor and recipient in media not supplemented with amino acid (Figure 3A). These structures were absent in cocultures when the focal amino acid was externally supplied. Nanotubes were also absent in monocultures of donor and recipient strains indicating a role in amino acid exchange. This hypothesis was further verified by differentially labelling the cytoplasm of donor and recipient cells with plasmids that express either red or green fluorescent proteins. Quantifying the proportion of recipient cells that contained both cytoplasmic markers after 24 hours of growth in coculture using flow cytometry allowed us to determine the exchange of cytoplasmic materials between cells under our experimental conditions. Finding that all cocultures analysed comprised a significant proportion of auxotrophic cells containing both fluorescent proteins simultaneously confirmed that cytoplasmic materials such as protein and free amino acids have been transferred from donor to recipient cells (Figure 3B). However, it has been previously shown that the presence of the amino acid auxotrophic genotypes require for growth, prevents the formation of nanotubes (Pande, Shitut et al. 2015). Uncoupling the obligate dependency by supplementing the growth medium with saturating concentrations of the focal amino acid provided no evidence for a significant increase in double-labelled auxotrophs (Figure 3B), thus linking the establishment of these structures to the physiological requirement for amino acid cross-feeding.

**Figure 3.**
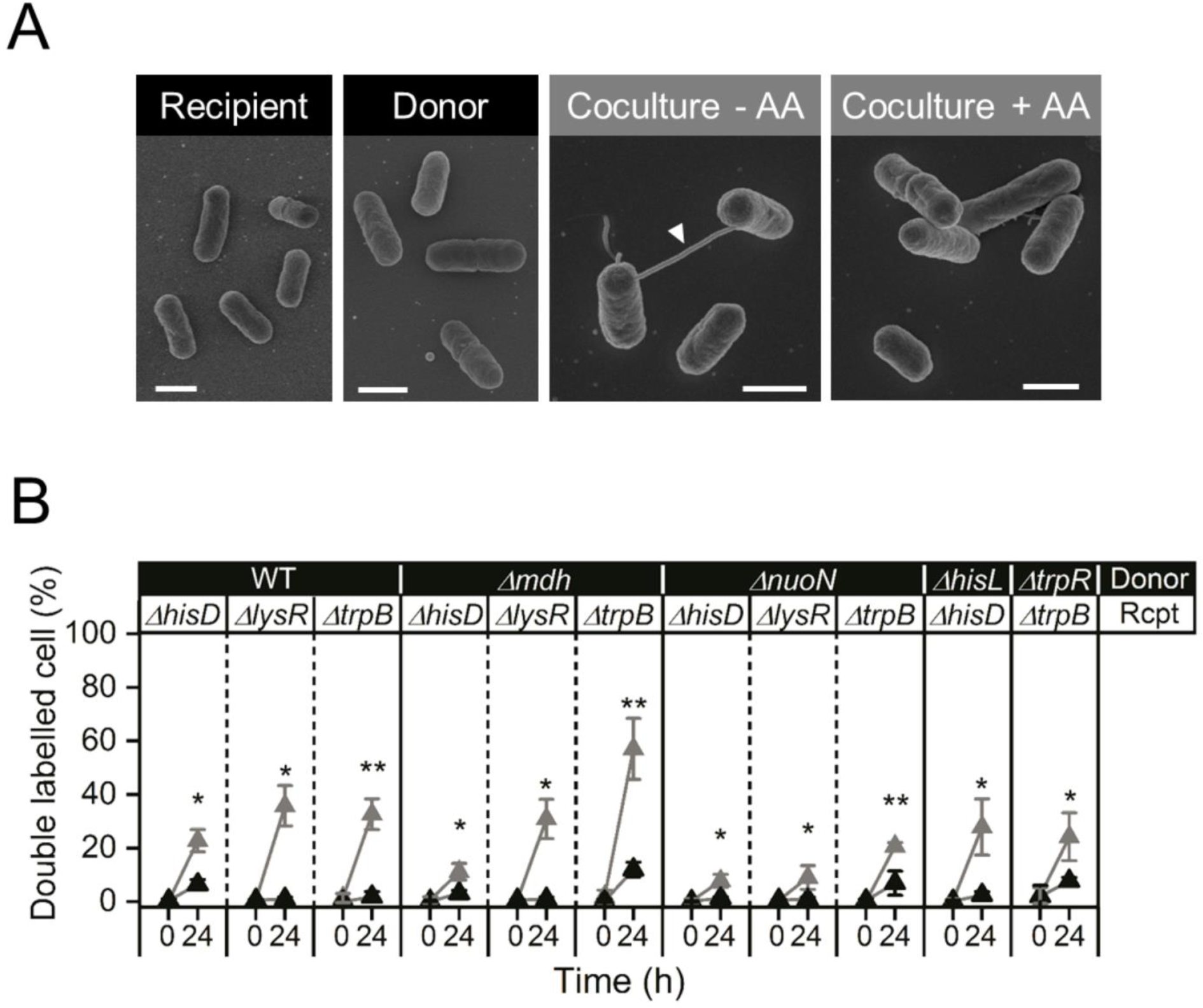
Exchange of cytoplasmic content through nanotubes. (**A**) Scanning EM images of recipient (Δ*hisD*) and donor (Δ*hisL*) in monoculture (black) and donor-recipient coculture (grey) conditions in the absence (− AA) and presence (+ AA) of amino acids after 24 hours of growth. Nanotubes (white pointer) were only observed in coculture without amino acid supplementation (Coculture - AA). Scale bars = 1 µm. (**B**) Cells exchange cytoplasmic material. The cytoplasm of donors and recipients were differentially labelled with the fluorescent proteins EGFP and mCherry, respectively. Quantifying the proportion of double-labelled auxotrophs containing both cytoplasmic markers after 0 h and 24 h of coculture allowed assessing an exchange of cytoplasm between bacterial cells. The experiment was conducted in the absence (grey triangles) and presence (black triangles) of the focal amino acid (100 µM). Asterisks indicate significant differences (paired t-test: ** P< 0.01, * P<0.05, n=4). In all cases, mean (±95% confidence interval) are shown.

### Auxotrophic recipients derive amino acid from cocultured donor cells

One hypothesis that could explain why recipients were able to grow in donor-recipient cocultures but not when cells were separated by a membrane filter (Figure 2A-C) is that the physical contact between cells increased amino acid production rates of donors. Amino acid production is energetically and metabolically very costly to the bacterial cell (Craig and Weber 1998, Akashi and Gojobori 2002, Kaleta, Schäuble et al. 2013). To minimize production costs, bacteria tightly regulate their amino acid biosynthesis, for example by end product-mediated feedback mechanisms that reduce production rates when cytoplasmic amino acid concentrations exceed critical thresholds (Thieffry, Huerta et al. 1998, Carlson 2007). In our case, recipient cells removed amino acids from the cytoplasm of donors using nanotubes. This decrease in the cell-internal amino acid pools could delay feedback inhibition in the donor cell, thus increasing its overall amino acid production (Figure 4). Quantifying the amount of free amino acids in the cytoplasm of donor cells in both the absence and presence of an auxotrophic recipient would allow testing the delayed-feedback inhibition hypothesis.

**Figure 4.**
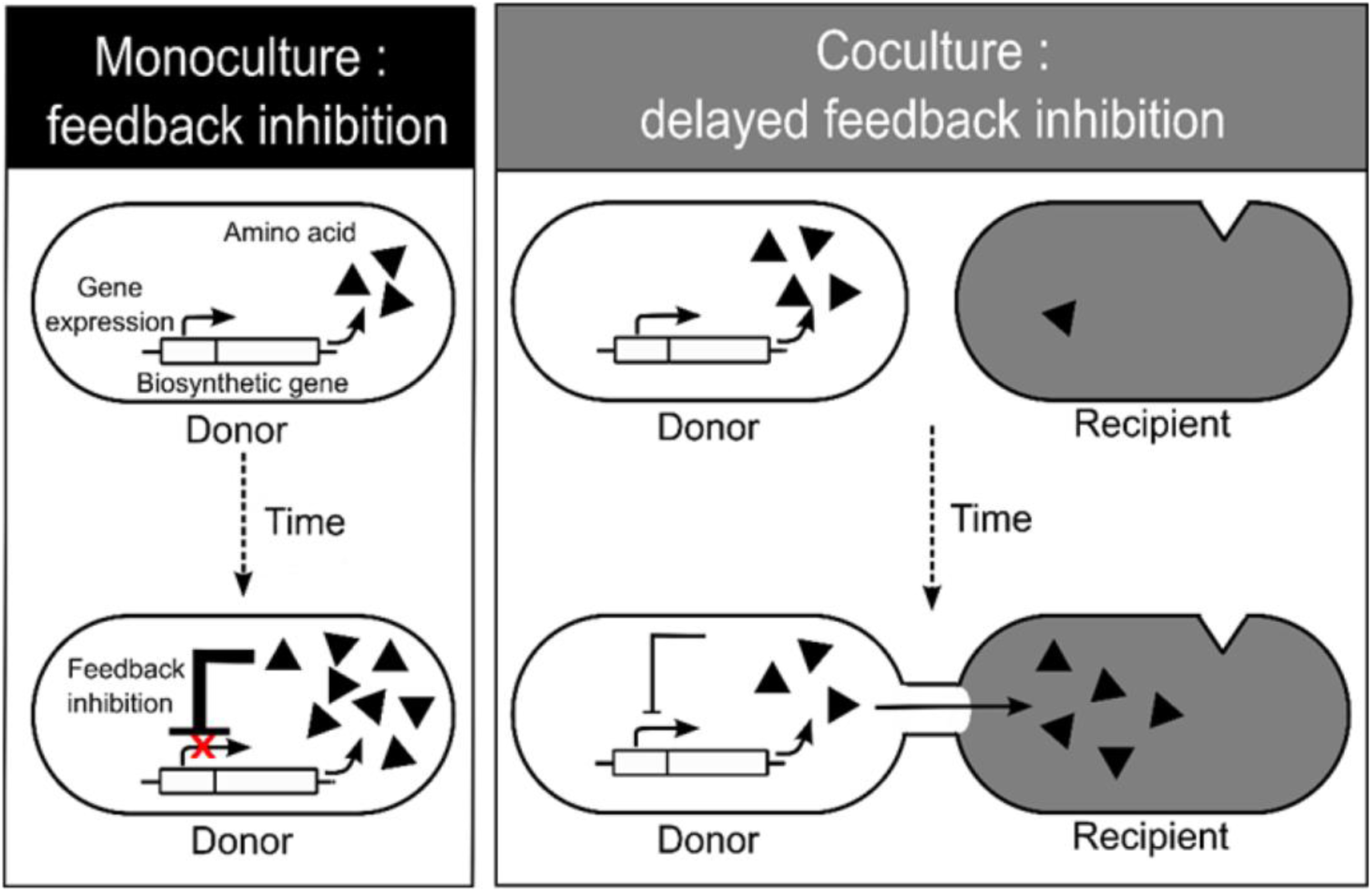
Delayed feedback inhibition hypothesis. In monoculture (left panel, black), amino acid concentrations in the cytoplasm of donor cells build up over time. When a certain concentration threshold is reached, these metabolites inhibit their own production by supressing the expression of the corresponding amino acid biosynthesis genes (i.e. end product-mediated feedback inhibition). In coculture (right panel, grey), auxotrophic recipients reduce cytoplasmic amino acid concentrations of donor cells. As a consequence, feedback inhibition of biosynthesis genes is delayed, thus resulting in an increased amino acid biosynthesis.

To determine cytoplasmic concentrations of free amino acids in real-time, we used the lysine riboswitch as a cell-internal biosensor. When free lysine binds to the riboswitch, it undergoes a conformational change, thus down-regulating expression of a downstream reporter gene, in our case *gfp* (Caron, Bastet et al. 2012). Introducing the plasmid-borne reporter construct (hereafter: *Lys-riboswitch*, Supplementary Figure 2) into the lysine auxotroph Δ*lysR* and exposing the resulting cells to different concentrations of lysine validated the utility of this biosensor: A strong negative correlation between the cells’ cytoplasmic amino acid concentrations as quantified via LC/MS/MS analysis of lysed cells and their fluorescence emission (r=-0.68, P=0.003, Supplementary Figure 3) corroborated that this construct allowed indeed determining levels of free lysine in the cytoplasm of living *E. coli* cells by simply quantifying their GFP emission.

Accordingly, introducing the lys-riboswitch into the lysine auxotrophic recipient (Δ*lysR*) and growing the resulting strain in lysine-supplemented media revealed consistently elevated levels of cytoplasmic lysine throughout the experiment (Figure 5B). In contrast, when the same recipient cells were grown in the absence of lysine, cell-internal lysine levels were significantly reduced (FDR-corrected paired sample t-tests: P<0.005, n=4, Figure 5B), indicating amino acid starvation of auxotrophic cells. Interestingly, when recipient cells were grown in the presence of one of the three donor genotypes, their lysine levels resembled that of lysine-starved auxotrophs until 18 hours of cocultivation, after which lysine levels increased back to the level of lysine-supplemented cells (FDR-corrected paired sample t-tests: P<0.04, n=4, Figure 5B). Prior to these coculture experiments, auxotrophs had to be pre-cultured in lysine-containing medium. Thus, the lysine levels measured in auxotrophs under coculture conditions likely reflected the fact that these cells first used up internal residual lysine pools before switching to other sources, in this case the cytoplasmic lysine of donor cells. Consistent with this interpretation is the observation that the presence of donor cells that provided this amino acid allowed lysine auxotrophs to grow (Figure 5A). A strongly positive correlation between the growth of lysine auxotrophs and their cell-internal lysine levels corroborates that the lysine auxotrophic recipients obtained from cocultured donor cells indeed limited their growth (r=0.625, P=0.003, Supplementary Figure 4).

**Figure 5.**
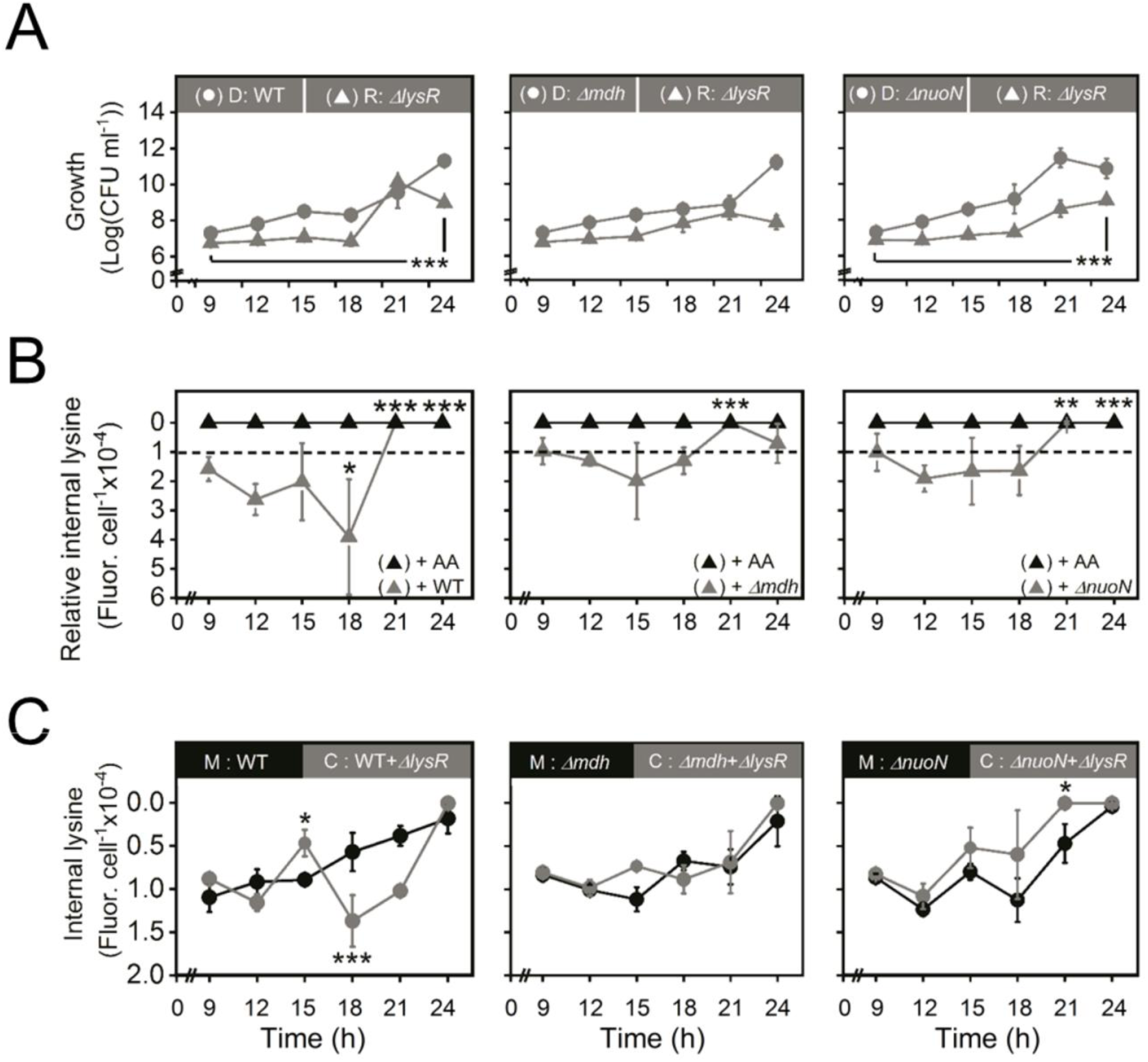
The presence of auxotrophs increases cytoplasmic amino acid levels in donor cells. (**A**) Growth of each partner in cocultures of donor (D, circles) and recipient (R, triangles) populations was determined as the number of colony-forming units (CFUs) ml^−1^ over 24 h. (**B, C**) Cytoplasmic lysine levels were quantified by measuring GFP fluorescence emission from a cell-internal reporter and normalized per cell containing the reporter. Low fluorescence levels indicate high lysine levels (note the inverted y-axes). (**B**) Lysine levels in lysine-supplemented monocultures (+ AA) and un-supplemented cocultures were measured relative to lysine-starved monocultures (dashed line). In the presence of lysine, monocultures of the recipient (black triangles) showed constantly increased cytoplasmic lysine levels. In coculture with the donor (grey triangles), lysine levels in the recipient first declined and then increased back to the level of the +AA condition. (**C**) In coculture with lysine-auxotrophic recipients, cytoplasmic lysine levels of WT donor cells were significantly increased at 15 h of growth and significantly decreased at 18 h of growth in coculture (C, grey circles) relative to monoculture conditions (M, black circles). However, in case of the overproducers Δ*mdh* and Δ*nuoN*, cell-internal lysine levels did not vary between mono- and coculture conditions. In all cases, mean (±95% confidence interval) are shown and asterisks indicate the results of FDR-corrected paired sample t-tests (*P<0.05, ***P<0.001, n=4).

### The presence of auxotrophic recipients increases cytoplasmic amino acid concentrations in donor cells

To test the delayed-feedback inhibition hypothesis, the lys-riboswitch was introduced into the three donors WT, Δ*mdh*, and Δ*nuoN*. Each of these donor genotypes were then grown in monoculture as well as in coculture with the lysine-auxotrophic strain Δ*lysR*. In these donor-recipient pairs only the donor contained the reporter plasmid.

The amino acid biosynthesis of WT cells is most stringently controlled, thus preventing accumulation of free lysine in its cytoplasm. In contrast, the cytoplasm of the Δ*nuoN* strain was characterized by generally increased amino acid levels (Supplementary Figure 1). Similarly, deletion of the malate dehydrogenase gene caused an accumulation of citric acid cycle intermediates and thus a dysregulated amino acid biosynthesis in the Δ*mdh* mutant (Pande, Merker et al. 2014). Hence, removing lysine from the cytoplasm by auxotrophs is expected to trigger the strongest response in WT cells in terms of how much cytoplasmic lysine levels are increased. In contrast, higher concentrations of lysine or its biochemical precursors in the cytoplasm of the Δ*mdh-* and the Δ*nuoN* strain may prevent a lowering of the lysine concentration below the critical threshold that triggers a further production.

We tested these predictions by monitoring changes in intracellular lysine levels of donor cells using the lys-riboswitch. In monocultures, lysine levels increased steadily over time (Figure 5C). This pattern, however, changed in the presence of the auxotrophic recipient. When *E. coli* WT cells were used as donor, their cytoplasmic lysine levels first increased significantly over the levels WT cells reached in monoculture (FDR-corrected paired sample t-tests: P<0.03, n=4, Figure 5C). After that lysine levels dropped significantly before increasing back to monoculture levels (Figure 5C). The observed fluctuations in the lysine levels of the donor’s cytoplasm are consistent with a nanotube-mediated cell attachment that is contingent on the nutritional status of the receiving cell. In contrast, when Δ*mdh* and Δ*nuoN* were cocultured as donor strains together with the auxotrophic recipient, their cytoplasmic lysine levels did not fluctuate as seen before in cocultures with WT (Figure 5C). While the cytoplasmic lysine levels of Δ*mdh* did not differ between mono- and coculture conditions, the Δ*nuoN* strain showed significantly increased lysine levels in its cytoplasm towards the end of the coculture experiment relative to Δ*nuoN* monocultures (FDR-corrected paired sample t-tests: P<0.03, n=4, Figure 5C). Thus, these observations are in line with the above expectations and confirm that an auxotroph-mediated removal of amino acids from the donor’s cytoplasm was sufficient to prompt an increased amino acid biosynthesis levels in donor cells. Conversely, lysine-auxotrophic recipients displayed significantly increased lysine levels when cocultured with one of the donor genotypes relative to lysine-starved monocultures (Figure 5B). Both observations together suggest a unidirectional transfer of amino acids from donor to recipient cells that resulted in an up-regulated amino acid biosynthesis in donors that depended on the presence of auxotrophic genotypes. Hence, these findings concur with the delayed-feedback inhibition hypothesis (Figure 4).

### The presence of auxotrophic recipients increases transcription of biosynthesis genes in donor cells

Bacterial cells use feedback inhibition to maintain homeostasis of certain metabolites in their cytoplasm. Once metabolite levels drop below a certain threshold, production levels are increased to allow optimal growth (Umbarger 1978, Scott, Gunderson et al. 2010). In the case of amino acid biosynthesis, the promoter elements that control transcription of biosynthetic pathways are frequently highly sensitive to intracellular levels of the synthesized amino acid (Thieffry, Huerta et al. 1998), thus enhancing transcription of the operon when the focal amino acid is scarce. As soon as amino acid concentrations reach optimal levels, further transcription is blocked enzymatically (Blasi, Bruni et al. 1973) or by direct binding of the amino acid to the operon (Yanofsky, Platt et al. 1981).

Taking advantage of this principle, we employed plasmid-borne promoter-GFP-fusion constructs to identify transcriptional changes in amino acid biosynthesis genes (Supplementary Figure 2). These reporter constructs have been previously shown to accurately measure promoter activity with a high temporal resolution (Zaslaver, Bren et al. 2006). For analysing the focal cross-feeding interactions, fusion constructs for *hisL* and *trpL* were selected, which respond to changes in the cytoplasmic concentration of histidine (Ames, Tsang et al. 1983) and tryptophan (Yanofsky, Platt et al. 1981, Merino, Jensen et al. 2008), respectively. Correlating GFP emission levels with the cytoplasmic concentration of the corresponding amino acid as quantified chemically via LC/MS/MS, revealed a significantly negative relationship for both histidine (r=-0.407, P<0.001, Supplementary Figure 3B) and tryptophan (r=-0.237, P=0.038, Supplementary Figure 3C), confirming the link between transcription of metabolic genes and the cytoplasmic concentration of the corresponding amino acids.

These promoter-GFP-fusion constructs were introduced into donor cells (i.e. WT, Δ*mdh*, Δ*hisL*, and Δ*trpR*), which were then cultivated for 24 hours in the absence or presence of the auxotrophic recipients Δ*hisD* or Δ*trpB*. In line with expectations, the presence of auxotrophic recipients strongly increased transcription of the corresponding biosynthetic genes in both WT and Δ*trpR* donor cells as compared to monocultures of donors (FDR-corrected paired t-tests: P<0.05, n=4, Figure 6). Interestingly, the histidine overproducing donor Δ*hisL*, which was characterized by cytoplasmic histidine levels that were significantly lower than the one observed in WT cells (Supplementary Figure 1B), had a four-fold higher promoter activity when paired with the auxotrophic recipient. Only the biosynthetic activity of the Δ*mdh* overproducer was unresponsive to the presence of histidine- and tryptophan-auxotrophic recipient.

**Figure 6.**
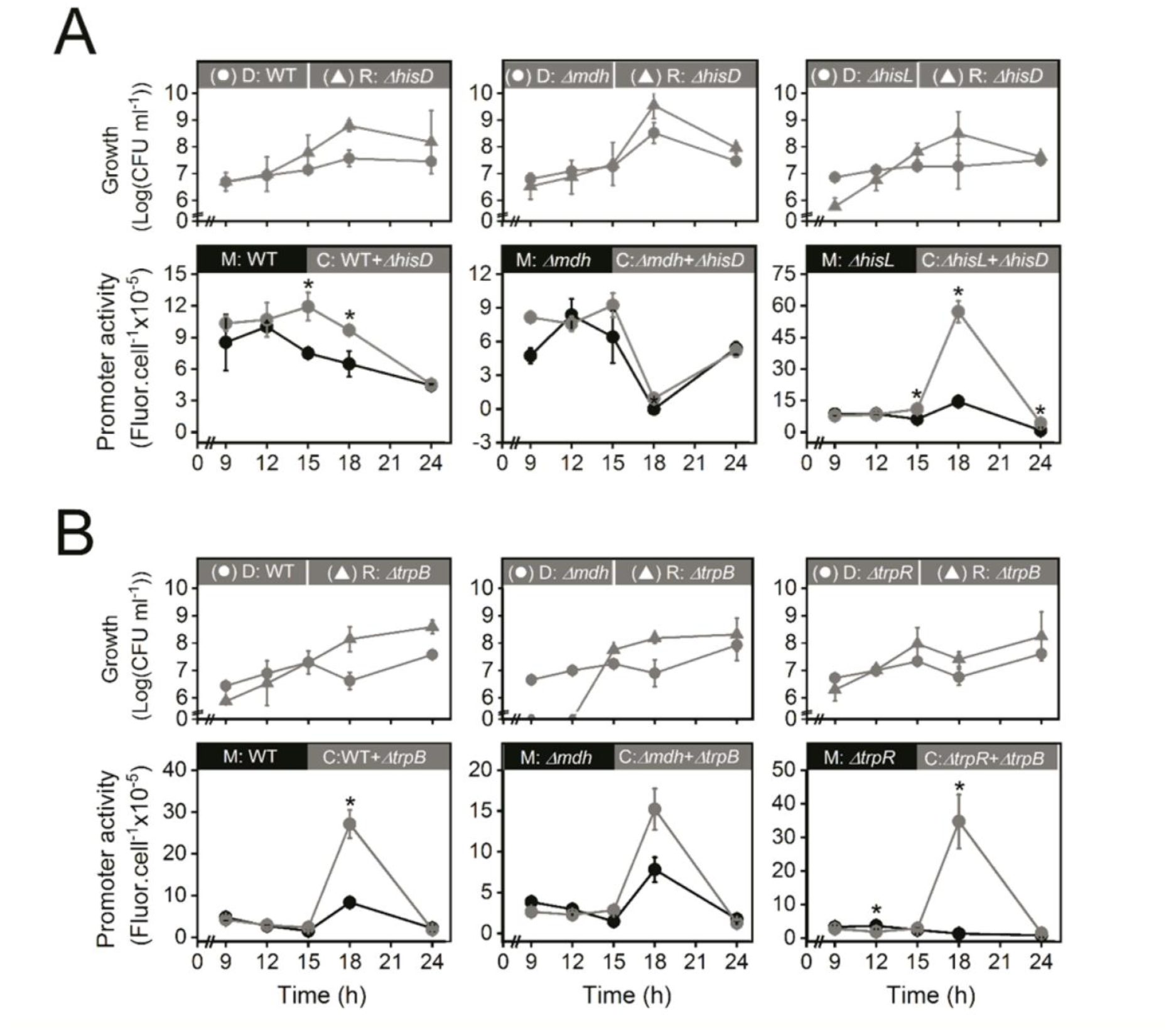
The presence of auxotrophs increases transcription of biosynthetic genes in the donor. (**A, B**) Top panels show growth of donor (D, circle) and recipient (R, triangle) in coculture over time quantified as the number of colony-forming units (CFU) per ml. Bottom panels show promoter activity of the donors’ amino acid biosynthesis gene in monoculture (M, black circles) and coculture with an auxotrophic recipient (C, grey circles). Promoter activity was quantified as the emission of GFP fluorescence from a promoter-GFP-fusion construct and normalized per number of donor cells (CFUs) containing the construct. Asterisks indicate significant differences of the promoter activity of donor cells in mono- and coculture conditions (FDR-corrected paired t-test: * P<0.05, n=4). Populations of donor cells (D, circles) were grown in monoculture or cultivated together with (**A**) the histidine auxotrophic recipient (Δ*hisD*) or (**B**) the tryptophan auxotrophic recipient (Δ*trpB*). In all cases, mean (±95% confidence interval) are shown.

Together, these results demonstrate that the presence of auxotrophic recipients significantly increased the amino acid production of donor cells. By withdrawing amino acids from the cytoplasm of donor cells, auxotrophic recipients prompted donor cells to readjust their amino acid levels by up-regulating the transcription of the corresponding amino acid biosynthesis genes.

## DISCUSSION

Our study demonstrates for the first time that the deletion of a single metabolic gene from a bacterial genome can be sufficient to physiologically couple the metabolism of two independent bacterial cells. Auxotrophic cells that had lost the ability to autonomously produce a certain amino acid established intercellular nanotubes to derive the amino acid they required for growth from other cells in the environment. Quantifying cell-internal amino acid levels as well as transcriptional activities of donor cells revealed an intercellular regulation of amino acid biosynthesis between donor and recipient in a source-sink-like manner. This relationship emerged as a consequence of feedback-based control mechanisms in the biosynthetic pathways of individual cells. These results show that a single loss-of-function mutation is not only sufficient to establish a metabolic interaction between two cells, but also that this can result in an intercellular regulation of amino acid biosynthesis that may help to reduce the costs arising from increased production levels.

The metabolic network of a cell provides and maintains specific levels of the building block metabolites that are required for growth (Holms 1996). An excess or deficit of metabolites within cells can disturb the cell-internal equilibrium and thus cause stress (Lee, Trostel et al. 2009, Grüning, Lehrach et al. 2010). For example, suboptimal metabolite concentrations in a cell’s cytoplasm can lead to osmotic imbalance (Csonka 1989) or cause oxidative stress (Chechik, Oh et al. 2008). To avoid these detrimental effects, bacteria have evolved complex mechanisms to sense and tightly control cytoplasmic metabolite levels. In the case of amino acids, this involves transcription attenuation, transcriptional repression, and feed-back enzyme inhibition (Umbarger 1978, Chopin 1993, Yanofsky 2004). If entering into a metabolic interaction with another organism negatively affected cellular homeostasis, this fact would represent a significant hurdle for the establishment of such interactions. However, finding that the same mechanisms that control metabolite production within cells also operate when some of the produced metabolites are transferred to other cells implies that consortia consisting of multiple cells may also optimally control metabolite production levels according to the needs of the cells involved.

When two complementary amino auxotrophic genotypes interact, the cross-feeding consortium can gain a fitness advantage of up to 20% relative to prototrophic cells, when both parties produce sufficient amounts of the metabolite their respective partner requires for growth (Pande, Merker et al. 2014). The results of our study can help to explain this fitness advantage: by selectively upregulating biosynthetic pathways that enhance growth of the consortium, cells only invest resources into those metabolites that help their current interaction partner to grow. In our experiments, the cost of producing increased amounts of amino acids to support the growth of another cell was only detectable for tryptophan on the level of the growth rate achieved (Fig. 2D). This observation is in line with previously published data, which indicates that of all three amino acids analysed in this study, tryptophan is the one that incurs the highest metabolic cost (Akashi and Gojobori 2002, Kaleta, Schäuble et al. 2013). Nevertheless, the cell densities all tested donor genotypes reached after growing for 24 h were independent of whether or not auxotrophic genotypes were present in the same environment (Fig. 2A-C), suggesting growing donor cells compensated for these costs. This can also explain the abovementioned strong fitness advantage experienced by cells engaging in reciprocal interactions: if each of two interacting cells slightly increase the production levels of the metabolite their respective partner requires for growth, both cells save the costs to produce another metabolite at all. In total, more resources are saved than invested, thus resulting in a net advantage of cross-feeding relative to metabolic autonomy.

Nutritional stress and starvation of bacterial cells, as is for example induced by auxotrophy-causing mutations, is known to trigger an aggregative lifestyle (Beloin, Valle et al. 2004, Benomar, Ranava et al. 2015). In many cases, this physical contact is followed by an exchange of cytoplasmic contents between interacting cells (Jahn, Gallenberger et al. 2008, Benomar, Ranava et al. 2015, Pande, Shitut et al. 2015). Structurally related intercellular connections are known to be involved in short- and long-distance communication in many multicellular organisms (Wegener 2001, Belting and Wittrup 2008). In both cases, networks of interacting cells are challenged with the question how to optimally distribute molecules within the interaction network. While the intercellular communication within tissues of eukaryotic organisms is notoriously difficult to study, our system provides a paradigmatic case to experimentally study the constraints and rules that determine the assembly and structure of intercellular communication networks. In this context, the results of our study indicate that the distribution of metabolites within networks of interacting bacterial cells may be self-organized by local interactions among neighbouring cells rather than requiring a superordinated regulatory system.

A functionally fused metabolism of two previously independent organisms as observed in this study, is strikingly reminiscent of the metabolic relationships that are frequently observed in symbiotic or pathogenic bacteria that infect eukaryotic hosts. For example, in the obligate association between aphids (*Acyrthosiphon pisum*) and their endosymbiotic bacteria *Buchnera aphidicola*, the aphid host regulates the amino acid production levels of its symbionts by changing its intracellular precursor concentrations (Russell, Poliakov et al. 2014). This functional link is afforded by a mutational elimination of feedback control in the corresponding biosynthetic pathway of the bacterial symbionts. Another case is the intracellular pathogen *Nematocida parisii* that secretes hexokinases to upregulate nucleotide biosynthesis of its host *Caenorhabditis elegans* (Dean, Hirt et al. 2016). Secretion of these enzymes results in increased levels of nucleotide biosynthetic precursors in the host cell, which the pathogen in turn utilizes for growth (Cuomo, Desjardins et al. 2012). Thus, by manipulating the biosynthetic pathway of their metabolite-producing partner, receiving individuals can ensure a sufficient supply with the required metabolite. Together, our study and the abovementioned examples exemplify, for metabolic interactions, how one partner can affect the biosynthesis of required metabolites of its counterpart by (i) increasing precursor levels (Dean, Hirt et al. 2016), (ii) enhancing the amount of key enzymes (Cuomo, Desjardins et al. 2012), or (iii) removing end products to delay feed-back inhibition (this study). Given that a metabolic complementarity on a genetic level is commonly observed in many symbiotic associations (Zientz, Dandekar et al. 2004) and free-living bacterial communities (Garcia, Buck et al. 2015), attempts to manipulate metabolite production levels of other interaction partners in favour of the acting individual are likely prevalent as well. If production and transport of traded metabolites is limiting the performance of the obligately interacting consortium as a whole, natural selection should act on optimizing these features to maximize fitness on a consortium-level.

Our work highlights the ease, with which newly emerged auxotrophic bacterial cells can derive the required nutrients from other cells in their environment. Contact-dependent exchange mechanisms that are induced upon nutritional stress facilitate the establishment of metabolic interactions as well as safeguard the transfer of cytoplasmic materials from one cell to another one (Dubey and Ben-Yehuda, Benomar, Ranava et al. 2015, Pande, Shitut et al. 2015). The intercellular regulation discovered in this study limits the amount of the traded metabolite that needs to be produced to meet the actual demand of the receiving cell. This mechanism should help to economize invested resources on a cell-level, thus allowing optimal growth of the interacting community.

Given that a loss of seemingly essential biosynthetic genes is very common in bacteria (D’Souza, Waschina et al. 2014), it is well conceivable how this type of reductive genome evolution can result in the formation of multicellular networks of metabolically interacting bacteria (Pande and Kost 2017). Once a biosynthetic gene is lost, the resulting auxotrophic genotype is more likely to lose additional genes than to regain the lost function via horizontal gene transfer (Puigbò, Lobkovsky et al. 2014). Because dividing metabolic labour in this way can be highly advantageous for the bacteria involved (Johnson, Goldschmidt et al. 2012, Pande, Merker et al. 2014), in their natural environment bacteria may exist within networks of multiple bacterial cells that reciprocally exchange essential metabolites.

## EXPERIMENTAL PROCEDURES

### Strains and plasmids used in the study

*Escherichia coli* BW25113 was used as wild type, from which mutants that overproduce amino acids (Δ*mdh*, Δ*nuoN*, Δ*hisL*, and Δ*trpR*) and mutants that are auxotrophic for histidine (Δ*hisD*), lysine (Δ*lysR*), or tryptophan (Δ*trpB*) were obtained by a one-step gene inactivation method (Pande, Merker et al. 2014, Pande, Shitut et al. 2015) (Supplementary Table 1). Deletion alleles were transferred from existing single gene deletion mutants (i.e. the Keio collection (Baba, Ara et al. 2006)) into *E. coli* BW25113 using phage P1. The cytoplasm of all donor and recipient strains was labelled by introducing one of the two plasmids pJBA24-*egfp* and pJBA24-*mCherry*. These plasmids constitutively express the ampicillin resistance gene (*bla*) as well as either the fluorescent protein EGFP (*egfp*) or mCherry (*mCherry*). Two reporter constructs were used: (i) lys-riboswitch (pZE21-GFPaav-Lys) for measuring internal amino acid levels (lysine) and (ii) promoter fusion plasmids (pUAA6-His and pUA66-Trp) for measuring the transcriptional activity of the promoters *hisL* and *trpL*, respectively (see supplemental experimental procedures for plasmid construction and characterization of reporter constructs).

### Culturing methods and general procedures

Minimal media for *Azospirillum brasiliense* (MMAB) (Vanstockem, Michiels et al. 1987) without biotin and with fructose (5 gl^−1^) instead of malate as a carbon source served as the growth media in all experiments. The required amino acids (histidine, lysine, and tryptophan) were supplemented individually at a concentration of 100 µM. Cultures were incubated at a temperature of 30 °C and shaken at 220 rpm for all experiments. All strains were precultured in replicates by picking single colonies from lysogeny broth (LB) (Bertani 1951) agar plates and incubated for 18 hours. The next morning, precultures were diluted to an optical density (OD) of 0.1 at 600 nm as determined by a Tecan Infinite F200 Pro platereader (Tecan Group Ltd, Switzerland). 10 µl of these precultures were inoculated into 1 ml of MMAB. In case of cocultures, donor and recipient were mixed in a 1:1 ratio by co-inoculating 5 µl of each diluted preculture. To cultivate strains containing the lys-riboswitch, ampicillin was added at a concentration of 100 µg ml^−1^ and kanamycin was added at 50 µg ml^−1^ in case of strains containing the promoter-GFP-fusion constructs. Anhydrotetracycline (aTc) (Biomol GmbH, Hamburg, Germany) was added at a concentration of 42 ng ml^−1^ to induce expression of the lys-riboswitch.

### Contact-dependent exchange of amino acids

To determine if physical contact between cells is required for an exchange of amino acids between donor and recipient cells, a previously described method was used (Pande, Shitut et al. 2015). In brief, each donor (i.e. WT, Δ*mdh*, Δ*nuoN*, Δ*hisL*, and Δ*trpR*) was individually paired with each recipient (i.e. Δ*hisD*, Δ*lysR*, and Δ*trpB*) and every combination was inoculated together into a Nurmikko cell that allows cultivation of both populations either together in the same compartment or separated by a membrane filter (0.22 µm, Pall Corporation, Michigan, USA). The filter allows passage of free amino acids in the medium, but prevents direct interaction between cells. After inoculating 4 ml of MMAB, the apparatus was incubated for 24 h. Bacterial growth after 24 h was determined as colony forming units (CFU) per ml culture volume by plating the serially-diluted culture on MMAB agar plates that did or did not contain ampicillin or kanamycin for selection. The increase in cell number was calculated as the logarithm of the difference between the CFU counts determined at the onset (0 h) of the experiment and after 24 h. Each donor-recipient combination was replicated 4-times for both experimental conditions (i.e. with and without filter).

### Relative fitness measurement

To quantify the effect of amino acid production on the fitness of donors, the growth of donor genotypes in terms of CFU per ml was calculated for mono- and coculture conditions at 4 time points in the exponential phase of growth (i.e. 14 h, 16 h, 18 h, and 20 h). Each donor (i.e. WT, Δ*mdh*, Δ*nuoN*, Δ*hisL*, and Δ*trpR*) was individually paired with one of each recipient (i.e. Δ*hisD*, Δ*lysR*, and Δ*trpB*) as well as grown in monoculture. Every combination was replicated six times. A regression line was fitted to the individual data points of a given condition (mono- or coculture) and the slope of this line was calculated to obtain the growth rate. The relative fitness of different donors was determined by dividing the growth rate each genotype achieved in coculture by the value of its respective monoculture.

### Scanning electron microscopy

Donor and recipient genotypes were either mono- or cocultured in 1 ml of liquid MMAB with and without amino acid supplementation for 24 h. 1 ml of culture was then fixed using a 2.5% glutaraldehyde solution prepared in a sodium cacodylate buffer (0.1 M, pH 7.0) for 1 h at room temperature. All fixed samples were allowed to sediment onto poly-L-lysine-coated glass coverslips (Sigma-Aldrich) for an additional 1 h time period. The glass coverslips were sputter-coated with a gold layer (25 nm) in a BAL-TEC SCD005 Sputter Coater (BAL-TEC, Lichtenstein). The gold-coated samples were visualized using a LEO 1530 Gemini field emission scanning electron microscope (Carl Zeiss, Jena) at 5 kV acceleration voltage and a working distance of 5 mm using an in-lens secondary electron detector.

### Flow cytometric analysis of cytoplasmic protein transfer

A previously established protocol was applied to identify a transfer of cytoplasmic material from donor to recipient genotypes (Pande, Shitut et al. 2015). For this, pairs of donor and recipient cells with differentially labeled cytoplasm (i.e. containing EGFP or mCherry) were co-inoculated into 1 ml MMAB. At the beginning of the experiment (0 h) and after 24 h of growth, the sample was analyzed in a Partec CyFlow Space flow cytometer (Partec, Germany). In the flow cytometer, cells were excited at 488 nm with a blue solid-state laser (20 mV) and at 561 nm with a yellow solid-state laser (100 mV). Green (*egfp*) and red (*mCherry*) fluorescence emission was detected at 536 nm and 610 nm, respectively. *E. coli* WT devoid of any plasmid was used as a non-fluorescent control. The number of single- and double-labeled cells in a population was quantified at both time points. Data analysis and acquisition was done using the FlowMax software (Partec GmbH, Germany). The experiment was conducted by coculturing eGFP-labelled donor with mCherry-labelled recipient gentoypes and *vice versa* in all possible combinations (i.e. each donor paired with each recipient, except in case of Δ*hisL* and Δ*trpR*, which were only paired with Δ*hisD* and Δ*trpB*, respectively) for 24 h. Each combination was replicated 4-times.

### Fluorescence measurement

The fluorescence levels of cells containing the lys-riboswitch or the promoter-GFP-fusion constructs were measured by transferring 200 µl of the culture into a black 96-microwell plate (Nunc, Denmark) and inserting the plate into a Tecan Infinite F200 Pro platereader (Tecan Group Ltd, Switzerland). The plate was shaken for 5 seconds prior to excitation at 488 nm followed by emission detection at 536 nm. Fluorescence values were always recorded together with a cognate control measurement. In case of the lys-riboswitch, the uninduced plasmid-containing culture served this purpose, while in case of the promoter fusion constructs, the promoter-less plasmid (pUA66) was used as control.

### Statistical analysis

Normal distribution of data was assessed using the Kolmogorov-Smirnov test and data was considered to be normally distributed when P > 0.05. Homogeneity of variances was determined using the Levene’s test and variances were considered homogenous if P > 0.05. One-way ANOVA followed by a Dunnett’s T3 post hoc test was used to compare growth differences in the contact-dependent growth analysis. Differences in the fluorescence emission levels of donor cells in the presence and absence of a recipient were assessed with paired sample t-tests. The same test was used to compare the number of recipient (Δ*lysR*) CFUs at the start and at the end of the coculture experiments to detect donor-enabled growth. The False Discovery Rate (FDR) procedure of Benjamini *et al*. (2006) was applied to correct P values after multiple testing. Pearson product moment correlation provided identification of the statistical relationship between cytoplasmic amino acid levels and fluorescence emission as well as between cytoplasmic lysine level and growth of the Δ*lysR* recipient.

## ACKNOWLEDGEMENTS

The authors thank Michael Reichelt for help with LC/MS/MS measurements, Uri Alon for providing the pZE21-GFPaav plasmid, Olin Silander for providing the promoter-GFP-fusion plasmids, Uwe Sauer for the promoter-less plasmid, Helge Weingart for supplying the pJBA24-*egfp* plasmid, and Wilhelm Boland for support. This manuscript benefitted greatly from discussions with the EEE-group, Martin Kaltenpoth and his Symbiosis-group, as well as Erika Kothe. This work was funded by grants from the Volkswagen Foundation, the Jena School for Microbial Communication as well as the DFG (SFB 944/2-2016).

## AUTHOR CONTRIBUTIONS

CK and SS conceived the study, SS, CK, and SP designed the study. SS performed all experiments. SS and CK interpreted and analyzed the data. TA generated some plasmids for the study. SS and CK wrote the manuscript, all authors amended the manuscript.

## REFERENCES

Akashi, H., & Gojobori, T. (2002). Metabolic efficiency and amino acid composition in the proteomes of Escherichia coli and Bacillus subtilis. Proc Natl Acad Sci U S A, 99(6), 3695–3700.

Ames, B. N., Tsang, T. H., Buck, M., & Christman, M. F. (1983). The leader mRNA of the histidine attenuator region resembles tRNAHis: possible general regulatory implications. Proc Natl Acad Sci U S A, 80(17), 5240–5242.

Baba T, Ara T, Hasegawa M, Takai Y, Okumura Y, Baba M et al (2006). Construction of *Escherichia coli* K-12 in-frame, single-gene knockout mutants: the Keio collection. Mol Sys Biol 2: 2006.0008–2006.0008.

Belenguer A, Duncan SH, Calder AG, Holtrop G, Louis P, Lobley GE et al (2006). Two routes of metabolic cross-feeding between *Bifidobacterium adolescentis* and butyrate-producing anaerobes from the human gut. Appl Environ Microbiol 72: 3593–3599.

Beloin C, Valle J, Latour-Lambert P, Faure P, Kzreminski M, Balestrino D et al (2004). Global impact of mature biofilm lifestyle on *Escherichia coli* K-12 gene expression. Mol Microbiol 51: 659–674.

Belting, M., & Wittrup, A. (2008). Nanotubes, exosomes, and nucleic acid-binding peptides provide novel mechanisms of intercellular communication in eukaryotic cells: implications in health and disease. J Cell Biol, 183(7), 1187–1191.

Benomar, S., Ranava, D., Cárdenas, M. L., Trably, E., Rafrafi, Y., Ducret, A.,… Giudici-Orticoni, M.-T. (2015). Nutritional stress induces exchange of cell material and energetic coupling between bacterial species. Nat Commun, 6: 6283.

Bertani G (1951). Studies on lysogenesis I.: The mode of phage liberation by lysogenic *Escherichia coli*. J Bacteriol 62: 293–300.

Bertels F, Merker H, Kost C (2012). Design and characterization of auxotrophy-based amino acid biosensors. PLoS One 7: e41349.

Blasi F, Bruni CB, Avitabile A, Deeley RG, Goldberger RF, Meyers MM (1973). Inhibition of transcription of the histidine operon *in vitro* by the first enzyme of the histidine pathway. Proc Natl Acad Sci U S A 70: 2692–2696.

Carlson RP (2007). Metabolic systems cost-benefit analysis for interpreting network structure and regulation. Bioinformatics 23: 1258–1264.

Caron MP, Bastet L, Lussier A, Simoneau-Roy M, Masse E, Lafontaine DA (2012). Dual-acting riboswitch control of translation initiation and mRNA decay. Proc Natl Acad Sci U S A 109: E3444–E3453.

Chechik G, Oh E, Rando O, Weissman J, Regev A, Koller D (2008). Activity motifs reveal principles of timing in transcriptional control of the yeast metabolic network. Nat Biotechnol 26: 1251.

Chopin A (1993). Organization and regulation of genes for amino acid biosynthesis in lactic acid bacteria. FEMS Microbiol rev 12: 21–37.

Craig CL, Weber RS (1998). Selection costs of amino acid substitutions in ColE1 and ColIa gene clusters harbored by *Escherichia coli*. Mol Biol Evol 15: 774–776.

Croft MT, Lawrence AD, Raux-Deery E, Warren MJ, Smith AG (2005). Algae acquire vitamin B12 through a symbiotic relationship with bacteria. Nature 438: 90.

Csonka LN (1989). Physiological and genetic responses of bacteria to osmotic stress. Microbiol Rev 53: 121–147.

Cuomo CA, Desjardins CA, Bakowski MA, Goldberg J, Ma AT, Becnel JJ et al (2012). Microsporidian genome analysis reveals evolutionary strategies for obligate intracellular growth. Genome Res 22: 2478–2488.

D’Souza G, Waschina S, Pande S, Bohl K, Kaleta C, Kost C (2014). Less is more: selective advantages can explain the prevalent loss of biosynthetic genes in bacteria. Evolution 68: 2559–2570.

D’Souza G, Kost C (2016). Experimental evolution of metabolic dependency in bacteria. PLoS Genet 12: e1006364.

Dean P, Hirt RP, Embley TM (2016). Microsporidia: why make nucleotides if you can steal them? PLoS Path 12: e1005870.

Dubey GP, Ben-Yehuda S Intercellular nanotubes mediate bacterial communication. Cell 144: 590–600.

Estrela S, Kerr B, Morris JJ (2016). Transitions in individuality through symbiosis. Curr Opin Microbiol 31: 191–198.

Garcia SL, Buck M, McMahon KD, Grossart H-P, Eiler A, Warnecke F (2015). Auxotrophy and intrapopulation complementary in the ‘interactome’ of a cultivated freshwater model community. Mol Ecol 24: 4449–4459.

Giovannoni SJ, Tripp HJ, Givan S, Podar M, Vergin KL, Baptista D et al (2005). Genome streamlining in a cosmopolitan oceanic bacterium. Science 309: 1242–1245.

Grüning N-M, Lehrach H, Ralser M (2010). Regulatory crosstalk of the metabolic network. TIBS 35: 220–227.

Holms H (1996). Flux analysis and control of the central metabolic pathways in *Escherichia coli*. Fems Microbiol Rev 19: 85–116.

Hubalek V, Buck M, Tan B, Foght J, Wendeberg A, Berry D et al (2017). Vitamin and amino acid auxotrophy in anaerobic consortia operating under methanogenic conditions. mSystems 2: e00038–17.

Jahn U, Gallenberger M, Paper W, Junglas B, Eisenreich W, Stetter KO et al (2008). Nanoarchaeum equitans and *Ignicoccus hospitalis*: new insights into a unique, intimate association of two Archaea. J Bacteriol 190: 1743–1750.

Johnson DR, Goldschmidt F, Lilja EE, Ackermann M (2012). Metabolic specialization and the assembly of microbial communities. ISME J 6: 1985–1991.

Junglas B, Briegel A, Burghardt T, Walther P, Wirth R, Huber H et al (2008). Ignicoccus hospitalis and Nanoarchaeum equitans: ultrastructure, cell-cell interaction, and 3D reconstruction from serial sections of freeze-substituted cells and by electron cryotomography. Arch Microbiol 190: 395–408.

Kaleta C, Schäuble S, Rinas U, Schuster S (2013). Metabolic costs of amino acid and protein production in *Escherichia coli*. Biotechnol J 8: 1105–1114.

Kiers ET, Rousseau RA, West SA, Denison RF (2003). Host sanctions and the legume-rhizobium mutualism. Nature 425: 78.

Kun Á, Papp B, Szathmáry E (2008). Computational identification of obligatorily autocatalytic replicators embedded in metabolic networks. Genome Biol 9: R51–R51.

Lee SJ, Trostel A, Le P, Harinarayanan R, FitzGerald PC, Adhya S (2009). Cellular stress created by intermediary metabolite imbalances. Proc Natl Acad Sci U S A 106: 19515–19520.

Loera-Muro A, Jacques M, Avelar-González FJ, Labrie J, Tremblay YDN, Oropeza-Navarro R et al (2016). Auxotrophic *Actinobacillus pleurpneumoniae* grows in multispecies biofilms without the need for nicotinamide-adenine dinucleotide (NAD) supplementation. BMC Microbiol 16: 128.

McFall-Ngai MJ (2014). The importance of microbes in animal development: lessons from the Squid-Vibrio symbiosis. Ann Rev Microbiol 68: 177–194.

Merino E, Jensen RA, Yanofsky C (2008). Evolution of bacterial trp operons and their regulation. Curr Opin Microbiol 11: 78–86.

Morris JJ, Lenski RE, Zinser ER (2012). The Black Queen Hypothesis: evolution of dependencies through adaptive gene loss. MBio 3: e00036–00012.

Morris JJ (2015). Black Queen evolution: the role of leakiness in structuring microbial communities. Trends Genetics 31: 475–482.

Paczia N, Nilgen A, Lehmann T, Gätgens J, Wiechert W, Noack S (2012). Extensive exometabolome analysis reveals extended overflow metabolism in various microorganisms. Microb Cell Factories 11: 122.

Pande S, Merker H, Bohl K, Reichelt M, Schuster S, de Figueiredo LF et al (2014). Fitness and stability of obligate cross-feeding interactions that emerge upon gene loss in bacteria. ISME J 8: 953–962.

Pande S, Kaftan F, Lang S, Svatos A, Germerodt S, Kost C (2015a). Privatization of cooperative benefits stabilizes mutualistic cross-feeding interactions in spatially structured environments. ISME J 10: 1413–1423.

Pande S, Shitut S, Freund L, Westermann M, Bertels F, Colesie C et al (2015b). Metabolic cross-feeding via intercellular nanotubes among bacteria. Nat Commun 6: 6238.

Pande S, Kost C (2017). Bacterial unculturability and the formation of intercellular metabolic networks. Trends Microbiol 25: 349–361.

Payne TMB, Rouatt JW, Lochhead AG (1957). The relationship between soil bacteria with simple nutritional requirements and those requiring amino acids. Can J Microbiol 3: 73–80.

Ponomarova O, Patil KR (2015). Metabolic interactions in microbial communities: untangling the Gordian knot. Curr Opin Microbiol 27: 37–44.

Puigbò P, Lobkovsky AE, Kristensen DM, Wolf YI, Koonin EV (2014). Genomes in turmoil: quantification of genome dynamics in prokaryote supergenomes. BMC Biology 12: 1–19.

Rodionova IA, Li X, Plymale AE, Motamedchaboki K, Konopka AE, Romine MF et al (2015). Genomic distribution of B-vitamin auxotrophy and uptake transporters in environmental bacteria from the Chloroflexi phylum. Environ Microbiol Rep 7: 204–210.

Russell CW, Poliakov A, Haribal M, Jander G, van Wijk KJ, Douglas AE (2014). Matching the supply of bacterial nutrients to the nutritional demand of the animal host. Proc R Soc Lond B Biol Sci 281: 7.

Sahu B, Ray MK (2008). Auxotrophy in natural isolate: minimal requirements for growth of the Antarctic psychrotrophic bacterium *Pseudomonas syringae* Lz4W. J Basic Microbiol 48: 38–47.

Scott M, Gunderson CW, Mateescu EM, Zhang Z, Hwa T (2010). Interdependence of cell growth and gene expression: origins and consequences. Science 330: 1099–1102.

Seth EC, Taga ME (2014). Nutrient cross-feeding in the microbial world. Front Microbiol 5: 350.

Shiio I, Ocirc, Tsuka S-I, Ocirc, Takahashi M (1962). Effect of biotin on the bacterial formation of glutamic acid I. Glutamate formation and cellular permeability of amino acids. J Biochem 51: 56–62.

Sieuwerts S, Molenaar D, van Hijum SAFT, Beerthuyzen M, Stevens MJA, Janssen PWM et al (2010). Mixed-Culture transcriptome analysis reveals the molecular basis of mixed-culture growth in *Streptococcus thermophilus* and *Lactobacillus bulgaricus*. Appl Environ Microbiol 76: 7775–7784.

Thieffry D, Huerta AM, Perez-Rueda E, Collado-Vides J (1998). From specific gene regulation to genomic networks: a global analysis of transcriptional regulation in *Escherichia coli*. Bioessays 20: 433–440.

Umbarger HE (1978). Amino acid biosynthesis and its regulation. Ann Rev Biochem 47: 533–606.

Vanstockem M, Michiels K, Vanderleyden J, Van Gool AP (1987). Transposon mutagenesis of *Azospirillum brasilense* and *Azospirillum lipoferum*: physical analysis of Tn5 and Tn5-Mob insertion mutants. Appl Environ Microbiol 53: 410–415.

Vogel KJ, Moran NA (2011). Sources of variation in dietary requirements in an obligate nutritional symbiosis. Proc R Soc Lond B Biol Sci 278: 115–121.

Wegener J (2001). Cell Junctions. eLS. John Wiley & Sons, Ltd.

Yanofsky C, Platt T, Crawford IP, Nichols BP, Christie GE, Horowitz H et al (1981). The complete nucleotide sequence of the tryptophan operon of *Escherichia coli*. Nucleic Acids Res 9: 6647–6668.

Yanofsky C (2004). The different roles of tryptophan transfer RNA in regulating trp operon expression in *E. coli* versus *B. subtilis*. Trends Genet 20: 367–374.

Zaslaver A, Bren A, Ronen M, Itzkovitz S, Kikoin I, Shavit S et al (2006). A comprehensive library of fluorescent transcriptional reporters for *Escherichia coli*. Nat Meth 3: 623–628.

Zelezniak A, Andrejev S, Ponomarova O, Mende DR, Bork P, Patil KR (2015). Metabolic dependencies drive species co-occurrence in diverse microbial communities. Proc Natl Acad Sci U S A 112: 6449–6454.

Zientz E, Dandekar T, Gross R (2004). Metabolic interdependence of obligate intracellular bacteria and their insect hosts. Microbiol Mol Biol Rev 68: 745–770.

